# Protein–protein interactions shape *trans*-regulatory impact of genetic variation on protein expression and complex traits

**DOI:** 10.1101/2024.10.02.616321

**Authors:** Jinghui Li, Yang I. Li, Xuanyao Liu

## Abstract

Most genetic variants influence complex traits by affecting gene regulation. Yet, despite comprehensive catalogs of molecular QTLs, linking trait-associated variants to biological functions remains difficult. In this study, we re-analyzed large maps of protein QTLs (pQTLs) to show that genes with *trans*-pQTLs but without *cis*-pQTLs are under strong selective constraints and are highly enriched in GWAS loci. We found that *trans*-pQTLs and their *trans* targets are highly enriched in interacting protein pairs, and *trans*-pQTLs in coding regions are significantly enriched at protein-protein interactions (PPI) interfaces. By leveraging existing PPI annotations for *trans*-pQTL mapping, we identified 26,028 *trans*-pQTLs influencing 1,061 PPI clusters. The *trans*-pQTLs of PPIs colocalized with 66% GWAS loci per trait on average for 50 complex traits, helping in many cases to link GWAS loci to cellular function. Finally, we identified *trans*-pQTL effects at multiple autoimmune GWAS loci that converge on the same PPIs, pinpointing protein complexes and signaling pathways that show promising therapeutic target potential.

## Introduction

Human genetic variation influences nearly all phenotypic traits. Yet, the exact mechanism by which most genetic variants impact traits remains difficult to determine. A major finding from genome-wide association studies (GWAS) is that 80–90% of trait-associated variants are non-coding^1,2^. Thus, most trait-associated variants are expected to have regulatory effects on nearby genes.

To map the effects of genetic variants on gene regulation, hundreds of studies, including those of large consortia, have mapped *cis*-expression quantitative trait loci (eQTLs) in diverse organs and cell-types. These studies have greatly improved our understanding of how genetic variants impact mRNA expression levels in *cis* through various mechanisms, including transcription factor binding at enhancers or promoters and post-transcriptional processes such as RNA processing^3–5^ or protein turnover^6^.

Even with these advances, for most trait-associated loci, it remains challenging to determine the mechanistic connection between the implicated genes and the trait in question, possibly because the genes play an indirect or peripheral role. We previously suggested that *trans*-regulatory effects can connect these peripheral genes to genes with direct trait effects^7^. Therefore, mapping *trans*-regulatory effects of genetic variants can help link GWAS signals to biological mechanisms^7,8^. Multiple studies now support this prediction. For example, the eQTLGen study^9^ detected a substantial number of *trans*-eQTLs and found that a third of GWAS trait-associated variants have clear *trans*-regulatory effects. These *trans*-eQTLs helped to pinpoint biological pathways associated with multiple immune diseases. Another study used Perturb-seq to infer gene *trans*-regulatory effects in endothelial cells, and identified multiple biological pathways showing convergence of genetic effects associated with coronary artery disease^10^. These studies, among others^11,12^, demonstrate the utility of mapping *trans*-regulatory effects to elucidate trait and disease mechanisms.

We recently developed a method, *trans*-PCO, to improve the mapping and interpretability of *trans*-eQTL effects^8^. Our work was inspired by the observation that although *trans*-eQTL effects are weak, they are often numerous and impact multiple co-expressed genes. We leveraged this observation by applying a multivariate approach to find the association of genetic variants with gene co-expression modules. Not only did this approach greatly increase *trans*-eQTL mapping power, it also helped us map the impact of many immune trait-associated variants on gene regulatory networks and biological pathways^8^.

While most studies have focused on *trans*-regulatory effects at the mRNA level, multiple large-scale proteomic studies have recently been conducted in plasma^13–16^. Technological advances in proteomics have enabled large protein expression studies that allow the detection and quantification of just a few thousand proteins, but with much larger sample sizes than that of existing *trans*-eQTL studies. Notably, the most recent UKB-PPP study^16^ measured plasma proteomic profiles of 54,219 participants, allowing the mapping of over fourteen thousand genetic associations with protein expression levels, including twelve thousand *trans*-pQTLs.

In this study, we sought to examine the *trans*-regulatory impact of genetic variants on the proteome and complex traits. To this end, we first carefully compared properties of pQTLs from UKB-PPP^16^ and eQTLs from multiple datasets^6,9^. Interestingly, we found that *trans*-pQTLs often have larger effect sizes than *trans*-eQTLs, and most *trans*-pQTLs are not *trans*-eQTLs, suggesting protein-specific *trans*-regulatory mechanisms. Indeed, we show that while *trans*-eQTLs are more likely to be mediated via transcriptional effects, a large number of *trans*-pQTLs affect protein expression level without impacting mRNA expression, implying a mechanism dependent on protein–protein interaction (PPI).

Additionally, we found that genes with *trans*-pQTLs but without *cis*-pQTLs are under stronger selective constraints than those with *cis*-pQTLs, and colocalize at a higher rate with GWAS loci than *cis*-pQTLs. These findings support a particularly important role for *trans*-pQTLs in mediating genetic effects on phenotypes. Leveraging *trans*-PCO and existing PPI databases, we identified over six thousand novel *trans*-regulatory associations between genetic variants and protein interaction networks. Many of these associations directly helped us link genetic variants to effects on protein complexes, which revealed biological functions impacted by trait-associated variants.

## Results

### *Trans*-effects are larger for protein than for mRNA expression levels

To study the impact of genetic variation on mRNA and protein expression levels in *trans*, we compared the effect sizes of eQTLs to that of pQTLs. To avoid winner’s curse, we first used pQTLs from UKB-PPP (plasma protein, sample size n = 52,363)^16^ to identify the *cis*- and *trans*-SNPs with the most extreme Z-scores for each protein and then used corresponding SNP-protein pairs from INTERVAL (plasma protein, n = 3,301)^13^ to ascertain the effect sizes of the top SNPs for each protein. We repeated this procedure for eQTLs, using (whole blood, n = 2,752)^17^ as the discovery dataset and DGN (whole blood, n = 913)^6^ for effect size ascertainment. We obtained 909 proteins and 11,322 genes that have corresponding *cis*- and *trans*-SNP–target pairs in both discovery and replication datasets. As expected, we found that for both eQTLs and pQTLs, *cis-* genetic variants had larger effects than *trans-* genetic variants, as measured by |Z-score|. However, while the effects of *cis*-eQTLs and *cis*-pQTLs are largely similar, *trans-*pQTL effect sizes were substantially larger than that of *trans*-eQTLs (**Figure 1a**). This trend is more evident when we compare the ratio of *trans*-to *cis*-Z-scores: this ratio for pQTLs were much higher than that for eQTLs (**Figure 1a**), implying *trans*-regulatory effect sizes were much larger for protein-level regulation than mRNA-level regulation.

**Figure 1.**
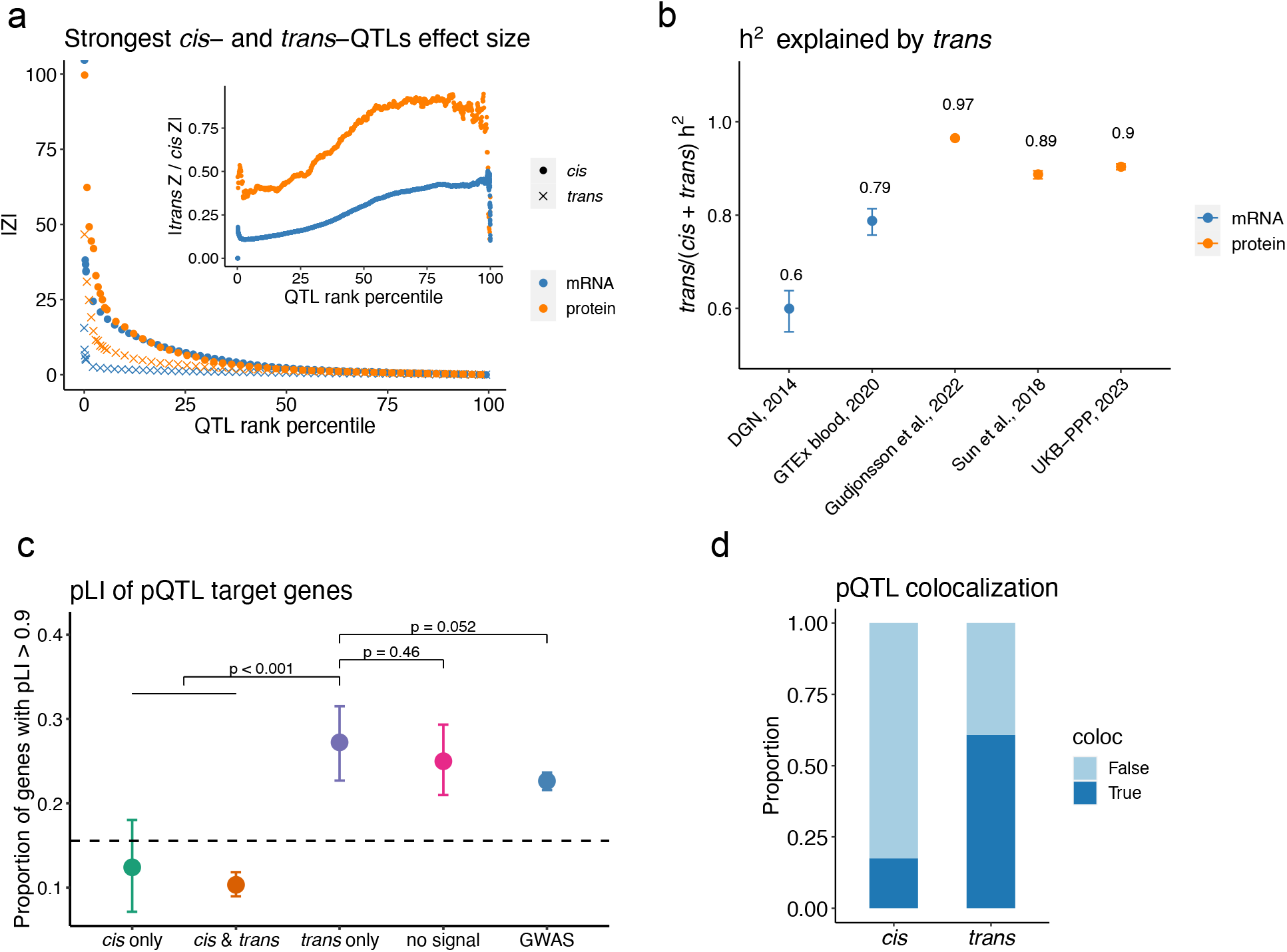
Properties of *trans*-pQTLs. a. The absolute z-score of the most significant *cis*- and *trans*-SNPs for each expressed protein in plasma and gene in whole blood. |Z| is sorted from largest to smallest for *cis*- and *trans*-SNP, respectively. The ratio of *trans* Z to *cis* Z of each rank is shown in the inset. b. Average proportion of heritability explained by *trans* for 662 common genes/proteins across 5 datasets. c. Proportion of pQTL target genes with pLI score > 0.9 in UKB-PPP and GWAS genes reported by ^19^. Dashed line shows the proportion of genes with pLI score > 0.9 across the whole genome. UKB-PPP genes are categorized into 4 groups: genes with both *cis*- and *trans*-pQTLs; with *cis*-pQTLs only; with *trans*-pQTLs only; and with no significant pQTLs. d. Proportions of *cis*- and *trans*-pQTLs colocalizing (PP4 > 0.75) with a GWAS locus for at least one trait out of 50 traits. Error bars in b and c show 95% confidence intervals determined by bootstrapping over 1,000 sampling iterations. P-values in c are calculated by a non-parametric test using the bootstrapping samples (**Methods**).

To further support these findings, we estimated the heritability of protein and mRNA expression explained by genetic variants, quantifying heritability explained by *cis*- and *trans*-variants separately for each gene^7,18^. On average, *trans*-genetic effects account 89%-97% of the heritability in protein expression level, while they account for 60%-79% of the heritability in mRNA expression levels (**Figure 1b**). The higher *trans-*heritability of protein expression is consistent with the relatively larger *trans*-effects on protein-level regulation.

### Genes with only *trans*-pQTLs are under stronger selection constraint than those with *cis*-pQTLs

Recent work has found that *cis*-eGenes are under weaker selective constraint than GWAS genes, suggesting that genes with *cis*-eQTLs are depleted among genes relevant to complex diseases^19^. Here, we ask whether genes with *trans*-pQTLs are similarly depleted. We classified *trans*-target genes (number of genes N = 2,837) into four different categories: (i) those with both *cis*-pQTL and *trans*-pQTL (N = 1,862), (ii) those with *cis*-pQTL but no *trans*-pQTL (N = 146), (iii) those with *trans*-pQTL but no *cis*-pQTL (N = 438), and (iv) those with neither *trans*-nor *cis*-pQTL (N = 391). We used the probability of being loss-of-function intolerant (pLI) as a proxy for selective constraint^19,20^. We found that genes with *cis*-pQTLs have lower proportions of high pLI-genes than “GWAS genes”, which are previously defined as genes that are near significant GWAS SNPs^19^ (10.8% vs 22.6%, p < 0.001), agreeing with previous observations (**Figure 1c**). Interestingly, we found that the proportion of high-pLI genes among genes with only *trans*-pQTLs (27.2%) was similar to that among GWAS genes (22.6%) and genes with no pQTLs (25.0%; p > 0.05), but was much higher than that among genes with *cis*-pQTLs (10.8%, p <0.001). We replicated these enrichment patterns using *trans*-pQTLs from two other proteomic datasets^13,21^ (**Figure S1**). These observations suggest that due to the strong selective constraints, although GWAS genes are depleted in strong *cis*-QTLs, they are more not depleted in *trans*-QTLs.

Next, we performed colocalization analyses between pQTLs and GWAS loci of 50 complex traits (**Table S1**) using coloc^22^. We found a higher proportion of *trans*-pQTLs than *cis-* pQTLs colocalizing with at least one GWAS locus (60.8% vs 17.5%, **Figure 1d**). Moreover, each *trans*-pQTL colocalizes with a mean of 4.1 complex traits, compared to a mean of 0.4 complex traits for *cis*-pQTL (**Figure S2**). Therefore, the limited overlap between *cis*-eQTLs and GWAS hits described previously^19,23^ does not appear to apply to *trans*-pQTLs. These findings strongly suggest that *trans*-pQTLs are more likely to influence complex traits and are also more pleiotropic (i.e., affect more traits) than *cis*-pQTLs.

In summary, given the strong selective constraints of pQTL *trans-*targets, and the large overlap between GWAS loci and *trans*-pQTLs, we conclude that mapping *trans*-regulatory effects may be particularly useful for studying and interpreting how GWAS variants impact complex traits.

### Protein–protein interaction is a major *trans*-pQTL mechanism

We have shown so far that *trans*-pQTL signals often have larger effect sizes than *trans*-eQTL signals, suggesting that there may be post-transcriptional *trans*-regulatory mechanisms influencing protein levels. Specifically, instead of impacting protein levels by first affecting mRNA levels of the *trans*-regulated genes, *trans*-pQTLs could directly influence protein synthesis or decay. Therefore, we first examined the sharing of *trans*-eQTLs and *trans*-pQTLs. Using pQTLs from the UKB-PPP dataset and eQTLs from two RNA-seq datasets in whole blood, DGN^6^ and eQTLGen^9^, we estimated the proportion of eQTLs that are pQTLs and, conversely, the proportion of pQTLs that are eQTL, by using the Storey π_1_ statistics^24^. We found high replication rates of *cis*-p/eQTLs with π_1_ ranging from 0.69 to 0.86 (**Figure 2a**). *Trans*-eQTLs also show high levels of replication in the UKB-PPP dataset (eQTLGen eQTLs π_1_ = 0.50; DGN eQTLs π_1_ = 0.77), confirming that *trans*-eQTLs often manifest as *trans*-pQTL, which is expected as changes in mRNA abundance generally result in a change in protein abundance. We thus take this as evidence that the UKB-PPP and whole blood RNA-seq datasets reliably capture the same genetic effects on gene regulation. Nonetheless, we found substantially lower rates of replication for *trans*-pQTLs among *trans*-eQTLs in both RNA-seq datasets (eQTLGen π_1_ = 0.24; DGN π_1_ = 0.11). Even when considering just the top 500 most significant *trans*-QTL signals as a way to account for a possible higher discovery power in the UKB-PPP dataset, *trans*-pQTLs were much less likely to be *trans*-eQTLs than the converse (π_1_ = 0.28 for *trans*-pQTLs in eQTLGen *trans*-eQTLs; π_1_ = 0.60 for eQTLGen *trans*-eQTLs in *trans*-pQTLs). These observations suggest that a major mechanism of *trans*-gene regulation involves direct effects on protein synthesis or decay, without impacting the transcription and/or mRNA levels of *trans*-target genes.

**Figure 2.**
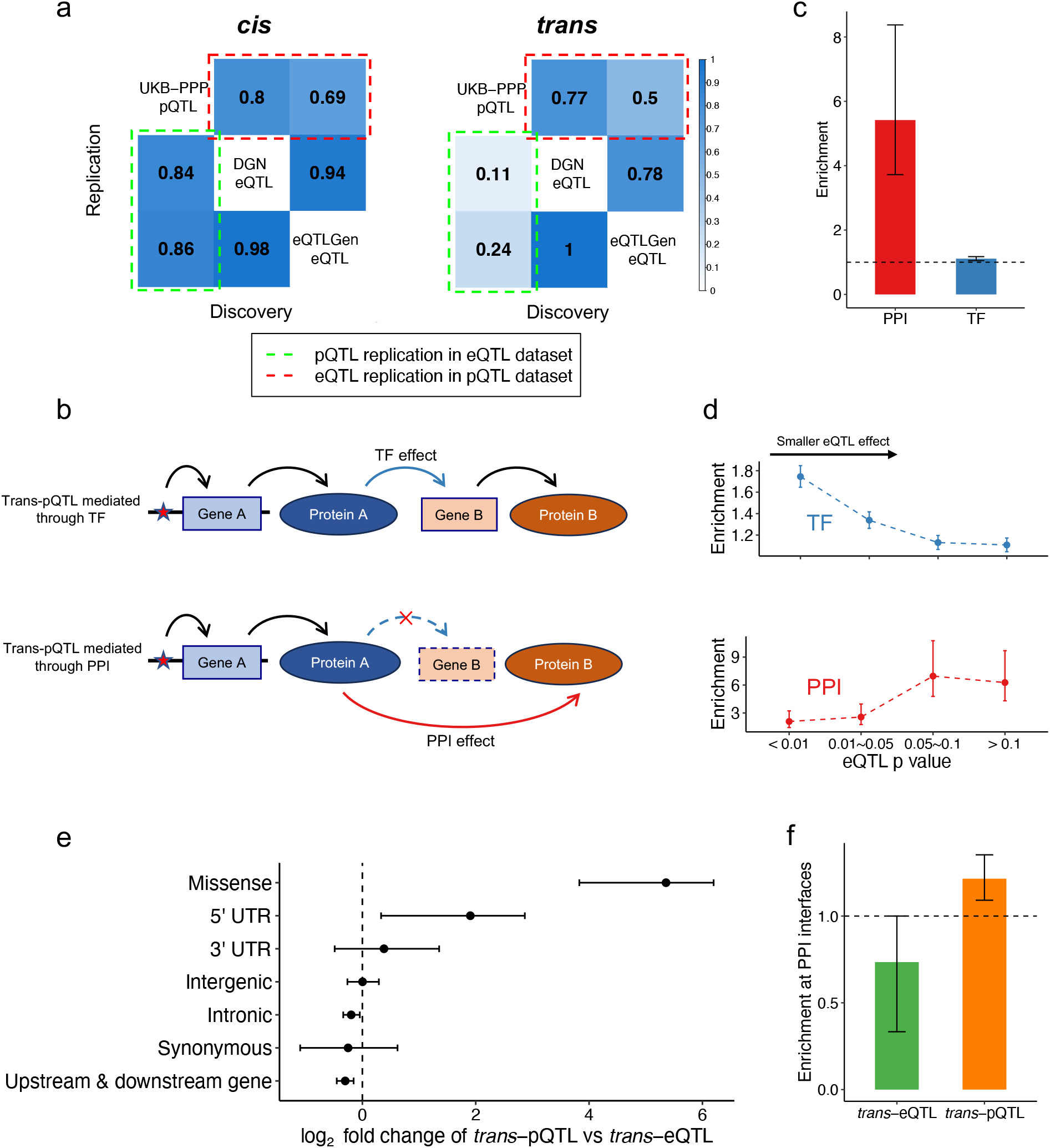
Protein-specific mechanism of *trans*-pQTLs. a.Estimated proportion of significant signals (π_1_) for *trans*-pQTLs/*trans*-eQTLs in gene/protein expression datasets. b. Probable pathways for *trans*-pQTLs affecting the target gene through a transcription factor (TF) mechanism and a protein–protein interaction (PPI) mechanism. c.Enrichment of nearest protein-coding genes of *trans*-pQTLs in TF and PPI (**Methods**). d.Enrichment of nearest protein-coding genes of *trans*-pQTLs in TF and PPI binned by different levels of eQTL p-values from DGN^6^. e. log_2_ fold change of the proportion of DGN *trans*-eQTLs and UKB-PPP *trans*-pQTLs for each SNP annotation decided by SnpEff^34^. f.Enrichment of DGN *trans*-eQTLs and UKB-PPP *trans*-pQTLs at PPI interfaces (**Methods**). PPI interfaces are from INCIDER^35^. Error bars in c to f show 95% confidence intervals determined by bootstrapping over 1,000 sampling iterations.

While plasma proteins contain circulating proteins derived from other tissue-types in addition to proteins from blood, we chose to compare UKB-PPP pQTLs to eQTLs in whole blood because 1) whole blood RNA-seq data has the largest sample sizes, and 2) we confirmed in GTEx data that UKB-PPP *cis*-pQTLs and *trans*-pQTLs have the highest replication rates in whole blood when compared to other tissues relevant to plasma proteins (i.e., liver or lymphoblastoid cell lines)^13^ (**Figure S3**).

Next, we predicted that unlike *trans*-eQTLs, for which a *cis*-effect on a transcription factor (TF) is a well-documented mechanism of action^8,9,25^, *trans*-pQTLs should be relatively depleted near TFs. We also hypothesized that protein–protein interactions (PPI), which involve physical binding and signaling pathways^26^ (Rao et al., 2014), may explain how *trans-*pQTLs function without any effects on mRNA levels (**Figure 2b**). Indeed, a seminal study on *trans*-pQTL in outbred mice found evidence that *cis*-e/pQTLs can affect protein expression levels in *trans* by altering protein-complex stoichiometry, leading to degradation of excess protein subunits^27^. To test the possibility that PPIs play a role in *trans*-regulation of protein expression levels in humans, we estimated the enrichment of *trans*-QTLs in interacting protein pairs (IPP), defined from 4 databases, including BioPlex^28^, CORUM^29^, HIPPIE^30^, and STRING^31^. We found a 5.4-fold enrichment (p-value < 0.001) of *trans*-pQTL regulator and target pairs in IPP, compared to random backgrounds (**Figure 2c**), suggesting *trans*-pQTL regulator are likely to be in the same PPI network as their *trans*-targets. In contrast, *trans*-pQTL regulators are only 1.1-fold enriched in TF (p-value < 0.001, **Figure 2c**) as compared to random backgrounds.

We next reasoned that *trans*-pQTLs with the most significant *trans*-eQTL p-values should represent *trans*-pQTLs that impact transcription and/or mRNA levels of the *trans*-target gene, while those with larger *trans*-eQTL p-values are more likely to function through PPI. To test this prediction, we binned *trans*-pQTL–protein pairs based on the association p-values of the corresponding *trans*-eQTL from the DGN dataset^6^ and estimated the enrichment of *trans*-pQTLs in IPP or TF. As predicted, we found that the enrichment in TF decreases with larger *trans*-eQTL p-values, whereas the enrichment in IPP increases with larger *trans*-eQTL p-values (**Figure 2d**). Altogether, these results indicate that interactions between proteins are a major protein-specific *trans*-regulatory mechanism.

### Coding variants of *trans*-pQTLs are enriched at PPI interfaces

Our previous work in *trans*-eQTLs showed that only 1% of *trans*-eQTL variants are in the coding region^8^. In contrast, we observed a surprisingly higher proportion, 14%, of *trans*-pQTL variants in coding regions. Comparing *trans*-pQTLs from UKB-PPP and *trans*-eQTLs from DGN, we found that *trans*-pQTLs have a significantly higher proportion of missense coding variants than *trans*-eQTLs (p < 0.001; **Figure 2e**). Moreover, *trans*-pQTLs that are coding variants have a 1.2-fold enrichment at PPI interfaces, compared to random backgrounds (p < 0.001; **Figure 2f**), while *trans*-eQTLs are depleted at PPI interfaces (p < 0.001; **Methods**). These results further highlight the differences between *trans*-eQTL and *trans*-pQTL and provide evidence of a potential *trans*-pQTL mechanism, namely the disruption of protein sequences at PPI interfaces^32^. As most *trans*-pQTLs are non-coding, we note that disruption of PPI interface is only one possible mechanism among many other plausible mechanisms, such as stoichiometry^27^ or co-translational mechanisms^33^.

### Large-scale mapping of *trans*-pQTLs using PPI annotations

Given the major role of PPIs in shaping the *trans*-effects of genetic variants on protein expression levels, we sought to use PPI annotations to improve the mapping and interpretation of *trans*-pQTLs. We recently developed a method, *trans*-PCO, which uses a multivariate association test to find *trans-*associations between SNPs and gene sets rather than single genes^8^. Compared to standard univariate tests used in the original DGN study^6^, *trans*-PCO substantially improves *trans*-eQTL detection power when a genetic variant impacts the expression level of multiple genes. Here, we use *trans*-PCO to improve mapping power of *trans*-pQTLs that impact the expression levels of multiple interacting proteins.

To define largely non-overlapping PPI clusters, we used the Markov CLuster algorithm (MCL)^36^ to separately cluster interacting proteins from BioPlex^28^, HIPPIE^30^, and STRING^31^ into PPI clusters. We used the protein complexes annotated in the CORUM database as is^29^. We then merged all PPI clusters together, which resulted in a final set of 1,088 PPI clusters encompassing 2,165 unique proteins assayed in UKB-PPP after removing overlaps (**Table S2**). The number of proteins per cluster ranges from 2 to 46 (mean = 3.47, median = 2). About half of the proteins (1,167) appeared in just one PPI cluster, while the remaining proteins appeared in multiple clusters as they formed different PPI clusters in different databases or were annotated as part of multiple CORUM protein complexes (**Figure S4**).

We performed genome-wide scans of *trans*-pQTLs of the 1,088 PPI clusters using *trans*-PCO, and identified 26,028 *trans*-pQTLs for 1,061 clusters at a Bonferroni-corrected threshold of p < 4.7×10^−11^ (**Figure 3a, Table S3**). These pQTL–PPI cluster pairs correspond to 1,535 independent *trans*-pQTL loci, each affecting a median of 4 PPIs (**Figure S5**). Interestingly, we observed 25 *trans*-pQTL “hotspots” that each affected more than 100 PPI clusters (**Figure 3b**). Twelve hotspots have been previously identified as *trans*-eQTL hotspots (regulating more than 10 genes) in eQTLGen^9^ and DGN^6^, including those near *ARHGEF3, ABO*, and *ATXN2*. While it is possible that some of the remaining 13 pQTL-specific hotspots may also be eQTL hotspots in related tissues, such as liver^16^, we verified that none are *trans*-eQTLs in GTEx liver. One pQTL-specific hotspots, *APOH* (affecting 100 PPI clusters), is a component of plasma circulating lipoprotein that has been implicated in several biological processes such as blood coagulation and cholesterol metabolism^37,38^. The involvement of *APOH* in a *trans*-regulatory hotspot may indicate an interesting post-transcriptional mechanism. We next compared PPI *trans*-pQTLs detected by *trans*-PCO with *trans*-pQTLs reported by the original UKB-PPP study. We found that 24% (6,234/26,028) of the significant associations have univariate association p-values between the same locus and proteins in the PPI cluster that are all larger than the UKB-PPP p-value threshold of 1.7×10^−11^. We considered these associations novel *trans*-pQTLs, while the remaining 76% of associations were considered shared *trans*-pQTLs.

**Figure 3.**
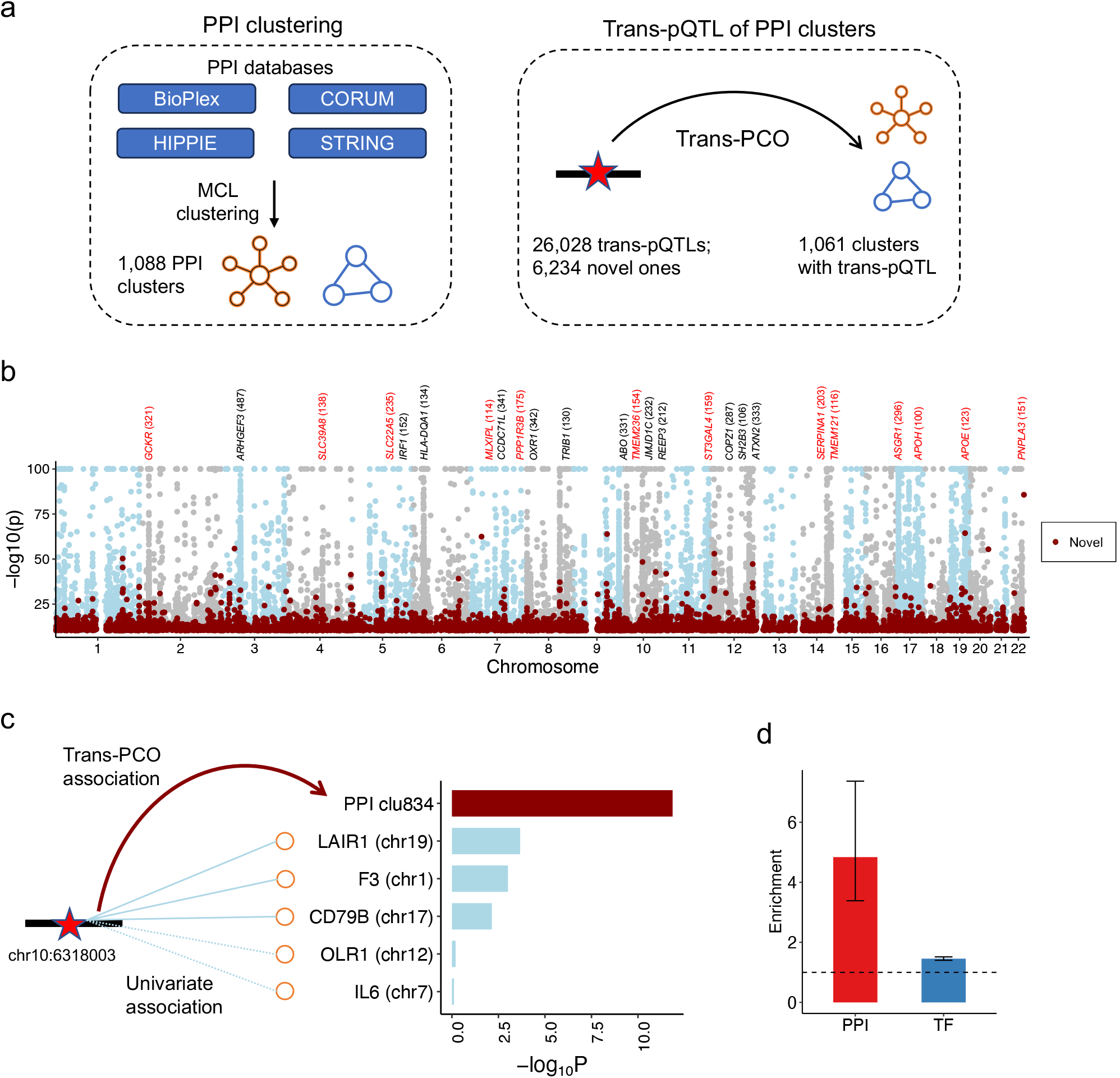
Detect *trans*-pQTLs using *trans*-PCO. a.Schematic of PPI clustering and the detection of *trans*-pQTLs of PPI clusters. b.Significant *trans*-pQTLs of PPI clusters detected by *trans*-PCO. The shared loci found by both *trans*-PCO and ^16^ are marked in blue and gray, while the novel loci not reported by ^16^ are marked in red. Twenty-five hotspots regulating more than 100 PPIs are marked above the figure, and those that are not *trans*-eQTL hotspots^6,9^ are marked in red. c.Example of a novel *trans*-pQTL identified by *trans*-PCO having multiple medium effects on protein subunits of the target PPI. d. Enrichment of nearest genes of novel *trans*-PCO pQTLs in transcription factors (TF) and PPI. Error bars show 95% confidence intervals determined by bootstrapping over 1,000 sampling iterations.

*Trans*-PCO identifies more *trans* signals by aggregating multiple weak effects. For example, a novel pQTL is significantly associated (p = 4.6×10^−11^) with a PPI cluster of five proteins (LAIR1, F3, CD79B, OLR1, and IL6), while each univariate association is far weaker (**Figure 3c**). Furthermore, to test if the PPI clusters are meaningful, we compared the MCL PPIs and random protein clusters. We found that PPI clusters not only have a larger correlation of univariate association z-scores than random clusters (0.19 vs 0.11, Mann-Whitney p = 5.9×10^−14^), but also more novel *trans*-pQTLs (Mann-Whitney p < 0.05; **Figure S6**). These results suggest that the protein clusters generated by MCL reflect true interactions among the proteins. Additionally, consistent with our findings that *trans-*pQTLs are more enriched in PPIs than TF activities, the novel *trans*-pQTLs have a 4.8-fold enrichment (p < 0.001) in IPP and only a slight enrichment in TFs (1.5-fold, p < 0.001; **Figure 3d**).

We found that novel PPI *trans*-pQTLs tend to affect a large proportion of proteins in the PPI rather than a single protein. We considered a *trans*-pQTL to affect multiple proteins in the target PPI when: 1) more than three proteins within the PPI have a univariate p-value <0.05/|PPI| (|PPI| is the number of proteins in the PPI), or 2) two or three proteins have univariate p-values <0.05/|PPI| and the proportion of these proteins in the PPI is > 0.2. Using these criteria, we obtained a total of 11,099 *trans*-pQTLs affecting multiple proteins of the target PPI. Overall, 56% of novel *trans*-pQTLs exhibit multiple univariate associations, compared to 38% of shared *trans*-pQTLs (**Figure S7**).

To explore possible mechanisms by which *trans*-pQTLs impact expression levels of proteins in PPIs, we examined the effect directions of *trans*-pQTLs on proteins in the same PPI. We reasoned that when the directions of a *trans*-pQTL’s effect are concordant for proteins in a PPI, it is more likely that they reflect a single direct effect on the entire PPI, rather than multiple separate effects on members of the PPI. Remarkably, of the 11,099 *trans*-pQTLs affecting multiple proteins of the target PPI, 58% have the same direction of effect on all proteins, which is 1.5-fold significantly enriched than random backgrounds (p-value < 0.001). The enrichment (> 9-fold) is even stronger for PPIs with 5 or more proteins (**Figure S8**). For example, a *trans*-pQTL (chr12:111,884,608) affects 27 proteins of PPI cluster 953 (univariate p < 0.05/|PPI|), and all the effects are negative (**Figure S8**). This observation agrees with the possibility that many *trans*-pQTL function by disrupting protein complex stoichiometry^27^, e.g. the paucity of one protein subunit results in a stoichiometric imbalance that leads to the degradation of other unbound protein complex members. Taken together, these results confirm that our *trans*-PCO analysis identified *bona fide trans*-acting variants that likely impact protein expression levels through protein interactions.

### *Trans*-pQTLs of PPI clusters improve interpretation of GWAS loci

To study GWAS loci in terms of their impact on PPIs, we conducted colocalization analyses between PPI *trans*-pQTLs and 50 complex traits (14 red blood cell traits, 11 white blood cell traits, 4 platelet traits, 8 immune-related diseases, 5 cardiovascular traits, 4 anthropometric traits, and 4 central nervous system disorders; **Table S1**). On average, 66% (27% ∼ 88%) of the GWAS loci associated with the 50 complex traits colocalize with PPI *trans*-pQTLs (**Figure 4a**).

**Figure 4.**
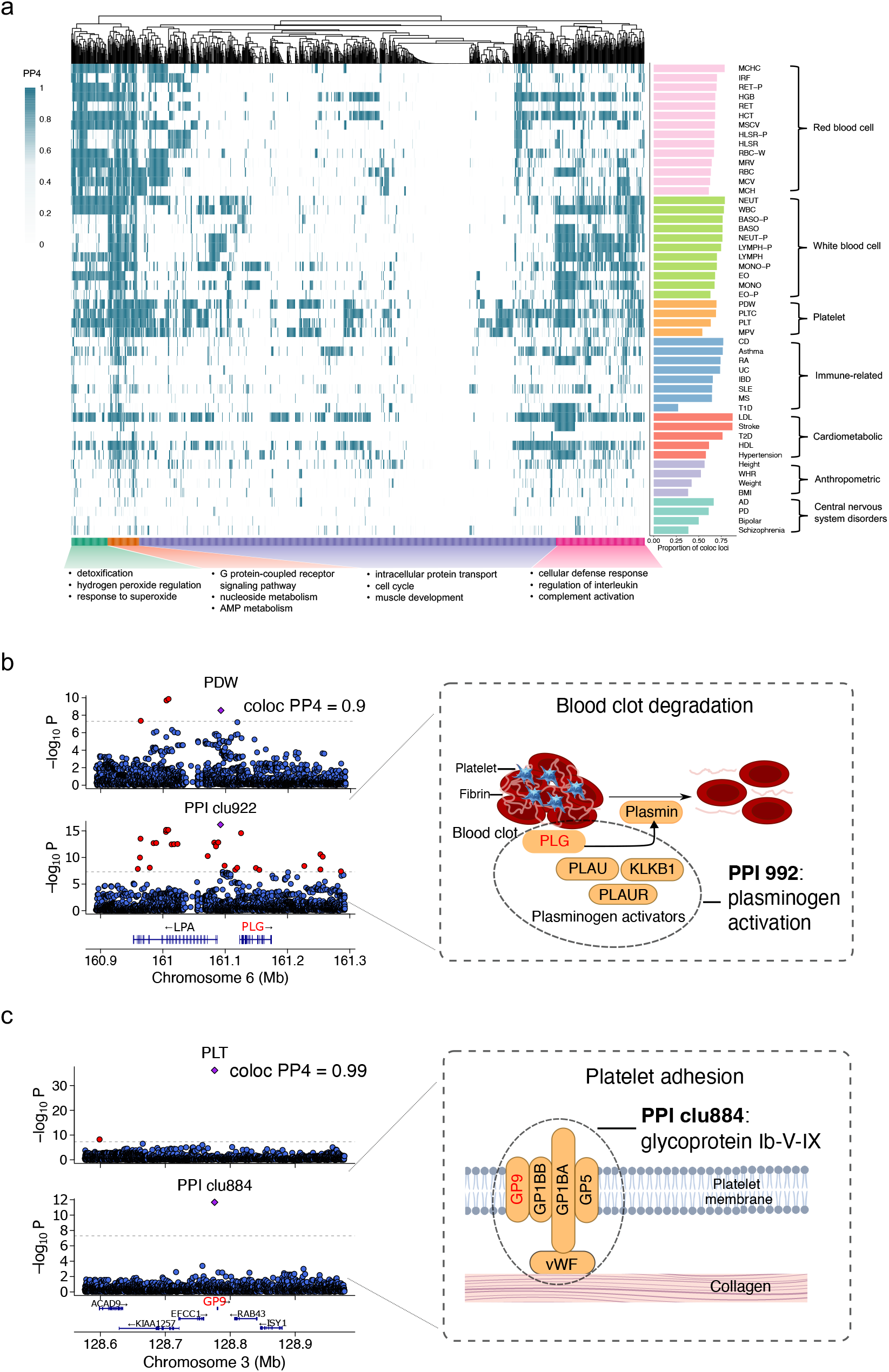
Colocalization between trans-pQTLs of PPIs and complex traits. a.The heatmap shows the most significant colocalization between each PPI and 50 traits. PPIs are divided into four groups based on their hierarchical clustering of the colocalization with each trait. The top three biological process GO terms that are unique to each group are listed below the heatmap. Proportions of GWAS loci colocalizing with trans-pQTLs of PPIs (PP4 > 0.75) for each trait are shown in the right panel. b. Colocalization between a *trans*-pQTL and a GWAS locus of platelet distribution width (PDW).c. Colocalization between a *trans*-pQTL and GWAS locus of platelet count (PLT).

Our analyses revealed that trait-associated loci have widespread *trans*-effects on the expression levels of interacting proteins. Nearly all (1,028/1,088) PPI clusters had at least one *trans*-pQTL colocalizing with at least one trait. We used hierarchical clustering to group traits and PPI clusters based on the highest colocalization (measured by PP4) between *trans*-pQTLs and GWAS loci from each of the 50 traits tested (**Figure 4a**). We divided the PPI clusters into 4 functionally related groups based on the colocalization clustering (**Table S4**) and analyzed the GO terms uniquely associated with each group (**Table S5 to S8**). The GO terms associated with the PPI group most likely to colocalize with white blood cell and immune-related trait hits included cellular defense (enrichment p-value = 1.4×10^−6^) and regulation of interleukin (enrichment p-value = 1.1×10^−5^). GO terms related to the group most likely to colocalize with red blood cell traits included detoxification (enrichment p-value = 1.6×10^−10^) and hydrogen peroxide metabolism (enrichment p-value = 4.0×10^−9^), highlighting the relevance of hemostasis regulation in blood.

We next showcase examples of colocalized PPI *trans*-pQTLs that directly help us interpret the functional impact of GWAS loci. We focused on colocalized *trans*-pQTLs affecting multiple proteins in the PPI, such that the GWAS loci are likely to function through the PPI. For example, we identified a GWAS locus associated with platelet distribution width (PDW) colocalizing with a *trans*-pQTL for PPI cluster 922 (PP4 = 0.90; **Figure 4b**), which is enriched in fibrinolysis regulation (enrichment p-value = 1.3×10^−13^). The colocalized locus is near *PLG*, which encodes plasminogen. Plasminogen and its activator proteins (PLAU, PLAUR and KLKB1) are all members of PPI cluster 992. The plasminogen activator proteins convert plasminogen to plasmin, which degrades fibrin and blood clots^39^. The colocalization signal supports the involvement of the PPI cluster functioning in plasminogen activation in PWD.

In another example, we found a colocalization between a locus associated with platelet count (PLT) and a *trans*-pQTL of PPI cluster 884 (PP4 = 0.99; **Figure 4c**), which is enriched in platelet activation (enrichment p = 4.1×10^−6^). The colocalized locus is near *GP9*, which encodes a glycoprotein on the platelet membrane^40^. GP9, along with other members of cluster 884 (GP1BB, GP1BA, GP5, and VWF), forms the glycoprotein ib-V-IX system, which is responsible for the platelet adhesion to collagens within the damaged connective tissues during the platelet activation process^41^. The colocalization supports that the PLT associated locus regulates the PPI cluster essential for platelet adhesion and activation.

These two examples, along with 1,610 other *trans*-pQTL colocalizations (**Table S9**), highlight *trans*-pQTLs that can impact a wide range of complex traits and diseases by impacting protein expression levels, many through protein–protein interactions.

### Convergence of multiple disease *trans*-pQTLs to PPIs

We have shown that *trans*-pQTL effects in PPIs can help with the functional interpretation of individual trait-associated GWAS loci. Further analyses of *trans-*pQTL can also help us prioritize PPIs that may be useful as therapeutic targets. Supporting this view, we found many instances where multiple GWAS loci of the same autoimmune disorder colocalize with *trans*-pQTLs that affect the same PPI, indicating that the PPI is especially relevant to the disease. For example, six rheumatoid arthritis (RA)-associated GWAS loci colocalize with trans-pQTLs of PPI cluster 960, which consists of TNFRSF13B (BAFF/APRIL receptor), TNFRSF13C (BAFF receptor), TNFRSF17 (BAFF/APRIL receptor), TNFSF13 (also known as APRIL), and TNFSF13B (BAFF cytokine) (**Figure 5a**). The convergence of GWAS loci to the same PPI in *trans* strongly suggests the functional importance of the PPI in RA. Indeed, cluster 960 comprises cytokines and receptors of the BAFF and APRIL systems, which induce activation of many important transcription factors (e.g. NFAT, AP1, and NF-kappa-B) and act as a potent B cell activator^42^. Both BAFF and APRIL systems have been reported to play an important role in the development of autoimmunity, especially in systemic lupus erythematosus (SLE), by sustaining B cell activation^42–44^. Notably, although a rare variant in the *BAFF* gene, was associated with multiple sclerosis, RA and SLE^45,46^, larger GWAS from other studies have not been able to identify genes in the BAFF/APRIL system as significantly associated with autoimmune diseases. This highlights the broad utility of *trans*-pQTLs for mapping variants to the biological functions that are disrupted in disease.

**Figure 5.**
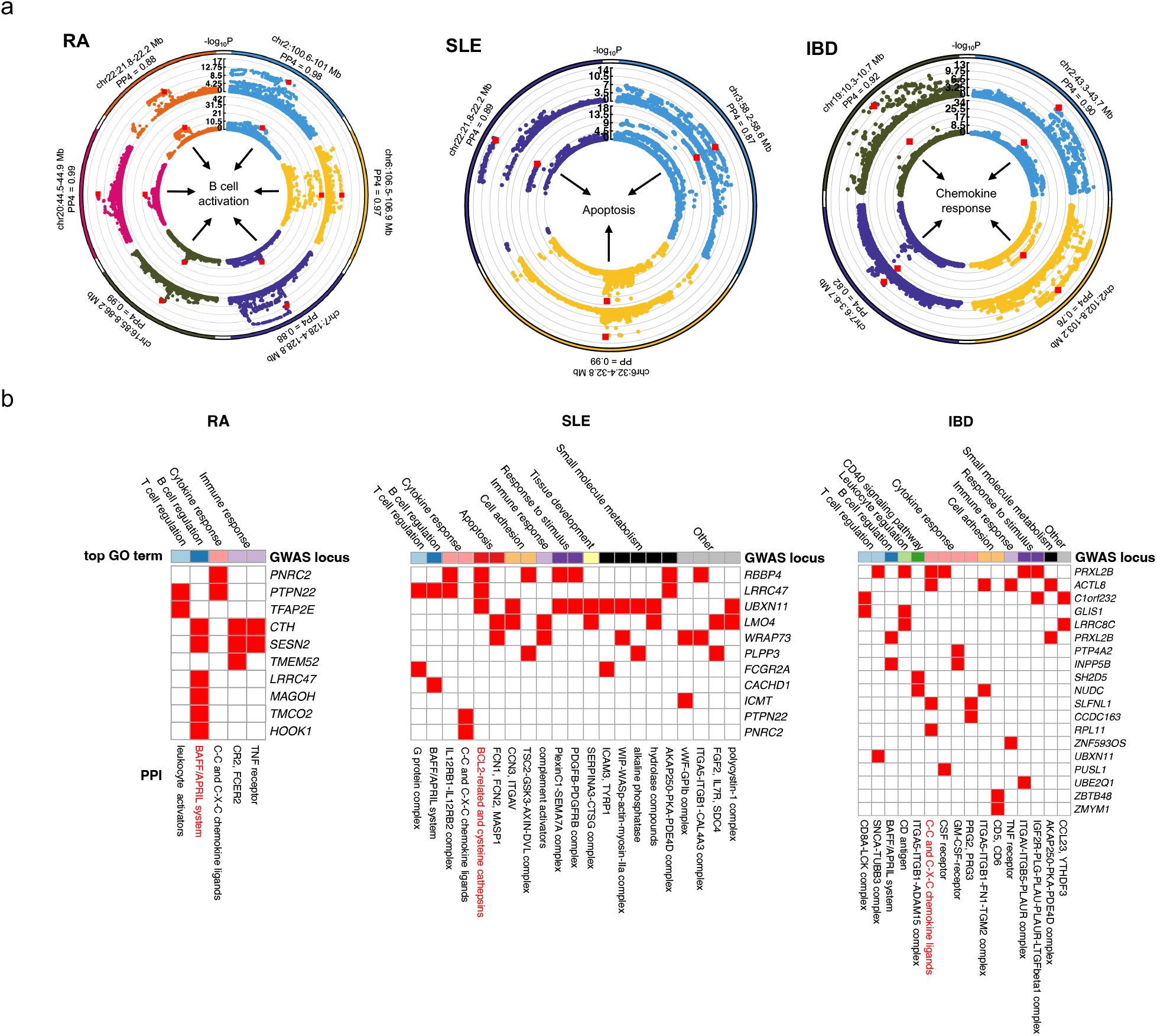
Multiple PPI *trans*-pQTLs converge to disease pathological pathways. a. The PPIs having the most colocalized GWAS/*trans*-pQTLs for RA, SLE and IBD. Each color represents a section of the genome in which a colocalization occurs, with the outer circle representing GWAS and the inner circle representing *trans*-pQTL. The leading SNP of each trans-pQTL is marked in red. The -log_10_ of p values for one locus (chr6:32.4 - 32.8 Mb) of *trans*-pQTL and SLE are shrunk by dividing by 10 for visualization.b. PPIs having multiple *trans*-pQTLs colocalizing with GWAS loci for RA, SLE, and IBD. Each colocalization between the GWAS locus and pQTL (PP4 > 0.75) is marked with a red block. The top biological process GO term enrichment of each PPI is annotated at the top of each panel.

In another example, three SLE-associated loci colocalize with *trans*-pQTLs of PPI cluster 1035 (**Figure 5a**), which is composed of BCL2-related proteins (BCL2, BCL2L1, BCL2L11, BID) and cysteine cathepsins proteins (CTSB, CTSC, CTSF, CTSH, CTSL, CTSO, CTSS, CTSV, and CTSZ). These proteins have been shown to regulate BCL2-dependent regulation of mitochondrial apoptotic pathways^47^. Additionally, cysteine cathepsins participate in antigen processing during immune responses, and impact cartilage degradation^48^. These processes play a role in several autoimmune diseases. Inhibition of cathepsin S has been proposed as a therapeutic modality for multiple autoimmune diseases^49^. Inhibition of CSTS was found to improve SLE symptoms by inhibiting autoantigen presentation in a mouse SLE model^50^, and also resulted in a significant reduction in disease in a mouse model of RA^49^. Other cathepsins such as CTSB, CTSC, and CTSL have also been proposed to be involved in autoimmune and inflammatory diseases^51–53^. Interestingly, in some contexts, BCL2 overexpression leads to a fatal SLE-like autoimmune disease^54^.

Finally, four inflammatory bowel disease (IBD) loci colocalized with *trans*-pQTLs for PPI cluster 964 (**Figure 5a**) composed of C-C motif chemokine ligands (CCL11, CCL13, CCL14, CCL17, CCL18, CCL2, CCL20, CCL21, CCL22, CCL23, CCL24, CCL25, CCL26, CCL28, CCL5, CCL8), C-X-C chemokine ligands (CXCL1, CXCL10, CXCL11, CXCL12, CXCL13, CXCL14, CXCL17, CXCL3, CXCL5, CXCL6, CXCL9, PF4) and two other proteins (PPBP, and XCL1). Chemokines are well known to be involved in a broad number of autoimmune diseases, including RA, SLE, and IBD^55–57^.

In total, we found 5, 21, and 16 PPIs with multiple significant GWAS locus/*trans-*pQTL colocalization events for RA, SLE, and IBD, respectively (**Figure 5b)**. These GWAS-converging PPIs (i.e., PPIs with multiple significant colocalization events in a single trait) are enriched in immune functions highly relevant to autoimmune diseases such as T cell and B cell regulation (enrichment p = 3.2×10^−44^), cytokine response (enrichment p = 1.8×10^−46^), and inflammatory responses (enrichment p = 1.4×10^−48^). The converged PPIs can be used to pinpoint critical pathological pathways that are missed in standard analyses of GWAS.

Furthermore, by analyzing converged PPIs that are shared across traits, we may be able to uncover shared genetic mechanisms between diseases. This raises the possibility of using PPIs identified here to elucidate drug targets or suggest the repositioning of existing drugs. For example, genetic loci associated with various autoimmune diseases exhibit convergence on both cluster 960 (BAFF/APRIL complex) and cluster 964 (C-C and C-X-C chemokines), which might indicate shared biological pathways across the diseases. Both Crohn’s disease (CD) and RA GWAS loci converge on cluster 960. Belimumab, a biological immunosuppressant that has been developed to treat SLE as a BAFF antagonist, was previously demonstrated to be efficacious and generally well tolerated in patients with RA^58^. Belimumab has been suggested as a potential treatment for IBD^59^ and, based on our results, may also be effective in CD. Additionally, the chemokine CCL11 in cluster 964 has been suggested as a drug target for asthma^60^. Our analyses suggest that CD and IBD GWAS loci also converge to impact this PPI, and patients with these conditions may also benefit from a drug targeting chemokine CCL11. Additional examples of convergence of disease-associated loci to PPI clusters are listed in **Table 1**.

**Table 1.**
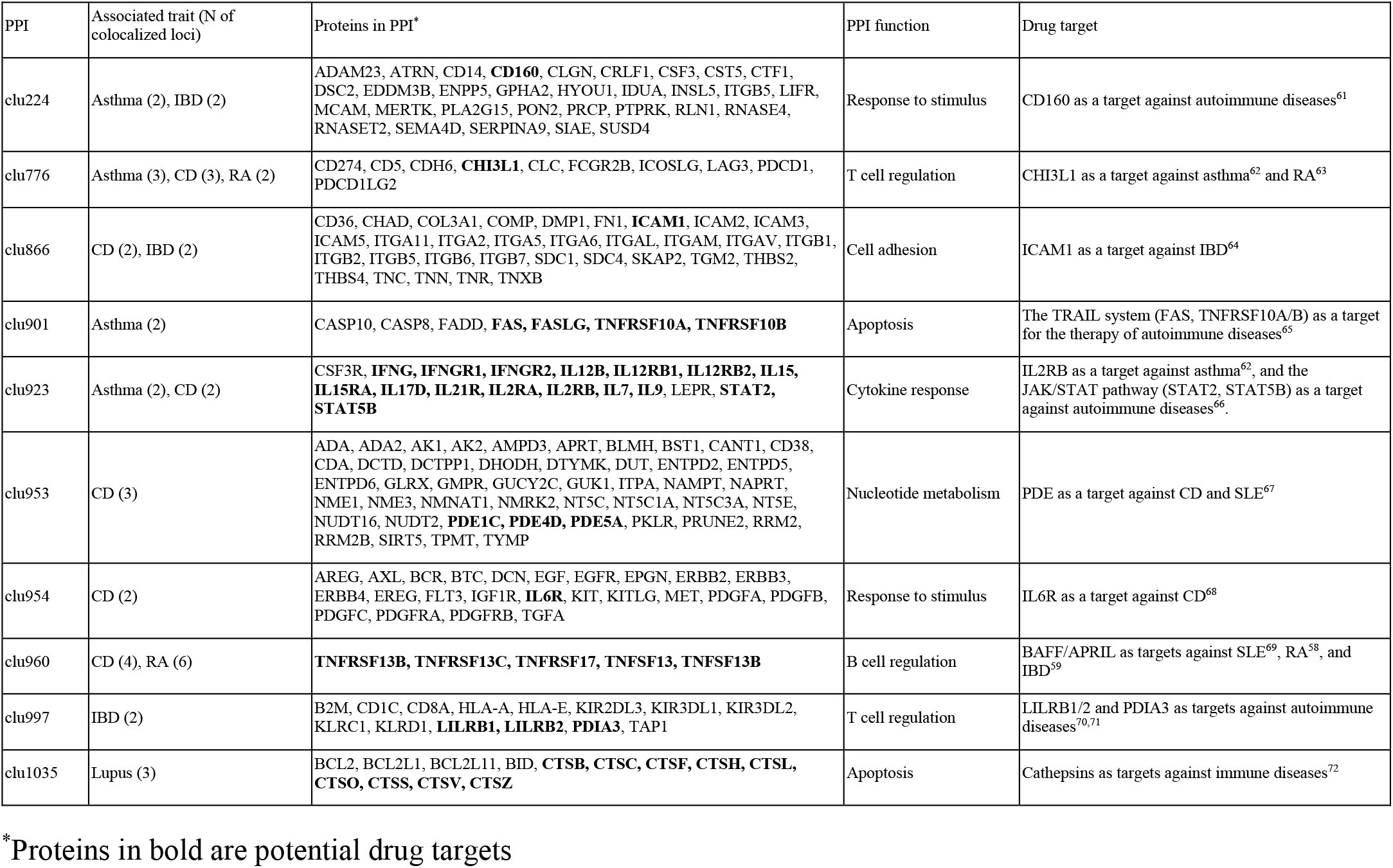
Drug target of proteins that are in disease-associated PPI clusters.

## Discussion

Most GWAS variants are non-coding, motivating a huge number of eQTL mapping studies with the goal of explaining the impact of non-coding variants on gene expression and cellular function. Nearly all eQTL mapping studies have thus far focused on mapping *cis*-gene regulatory effects exclusively. However, recent studies have found that steady state *cis*-eQTL studies are less likely to inform disease genetics, as genes with mapped *cis*-eQTLs show fundamental differences in several properties compared to disease genes^19^. In this study, we found that *trans*-pGenes (i.e., genes with at least one *trans-*pQTL) without any *cis*-pQTLs are under comparable selective constraint as disease genes, highlighting the importance of *trans*-QTLs in studying complex traits genetics. We also found that a large fraction of *trans*-pQTLs are not *trans*-eQTLs and have larger effect sizes than *trans*-eQTLs. Thus, they likely represent post-transcriptional mechanisms, such as those involving PPIs. In particular, we demonstrate that PPI is a major mediator of these post-transcriptional *trans*-effects.

We adapted *trans*-PCO, a method we previously developed to detect *trans-*eQTLs, to improve *trans*-pQTL mapping power by leveraging our finding that *trans-*pQTLs often act through PPIs. Using this approach, we mapped 26,028 *trans*-pQTLs of PPIs, which colocalize with over 60% of all GWAS loci analyzed. We show many cases where the colocalization of PPI *trans-* pQTLs and GWAS help us interpret how genetic effects percolate through PPIs to impact complex traits. We also highlighted several PPIs for which multiple GWAS loci converge to in *trans*. We speculate that these PPIs are not only important for understanding the genetic etiology of complex diseases, but they can also serve as potential drug targets.

There are several limitations of our study. We used the proteomic dataset from UKB-PPP^16^, which recruited more than 54,000 individuals. Despite the large sample size, it only includes less than 3,000 proteins. Given that the human body has ∼20,000 protein-coding genes and each has multiple transcript isoforms, the proteins considered in this study comprise a small proportion of the entire protein profile. Another limitation is that our association between *trans*-pQTLs and PPIs is dependent on existing maps of PPI networks. Currently, the knowledge of PPIs is limited and there is no comprehensive map of PPI networks. We have leveraged multiple PPI databases to construct PPI networks to mitigate the shortcomings of any one database. Additionally, it has been shown that PPIs may have cell-type specificity^28^, which is not considered in this study. Furthermore, we only investigated the role of PPI as a *trans*-pQTL mechanism. It is likely that other post-transcriptional regulation can also play a role in *trans* protein regulation, such as co-translational mechanisms^33^. We note that PPI is not unique to *trans*-pQTL; some *trans*-eQTLs may also change the expression of distal genes through PPI, though these are secondary effects that are weak. Lastly, the existence of *trans*-pQTL effects on multiple proteins in the PPI does not necessarily mean the *trans* effect through this mechanism. Further functional validation is required to draw definitive conclusions about the *trans*-effects on PPIs and their ultimate effects in diseases.

In conclusion, our study supports prevalent pQTL-specific *trans*-regulatory effects that depend on PPIs and play important roles in complex traits and diseases. Our work suggests that mapping *trans* regulatory-effects of trait-associated loci on the expression levels of proteins in PPI is a promising approach for inferring mechanisms of GWAS loci, and for identifying important protein drug targets.

## Methods

### Datasets

#### pQTL data

We used pQTL summary statistics from the UKB-PPP (n = 52,363) to generate the major results^16^. We use the same Bonferroni corrected p-value threshold (p < 1.7×10^−11^) as the UKB-PPP study to define significant associations. For each protein, we select the most significant SNP as the leading SNP and expand two flanking regions of size 250 kb around it (500 kb window). We exclude all the SNPs in the window and search for the next leading SNP and genomic window until all the significant SNPs are investigated. We merge overlapping windows into one and define it as a pQTL with the most significant SNP as the leading SNP. The major histocompatibility complex region (chr6: 28,477,797 to 33,448,354) is treated as one locus. We define a pQTL as *cis*-pQTL if the leading SNP is within 1 Mb from the transcription starting site (TSS) of the target gene, and as *trans*-pQTL if the leading SNP is more than 1 Mb from TSS or located on a different chromosome. The full list of pQTLs analyzed in this study is shown in **Table S10**. All the QTLs, including pQTLs and eQTLs, are defined using the same approach except with different p-value thresholds. All genomic coordinates are reported based on the GRCh37 assembly.

#### *h*^2^ of protein and mRNA expression

We applied stratified LD-score regression (v.1.0.1)^73^ to estimate the *cis* and *trans* heritability (*h*^2^) for protein and mRNA expression based on the summary statistics^18^ (Liu et al., 2017). In a stratified LD-score regression model, SNPs are partitioned into different functional categories. LD to a category contributing more to *h*^2^ will increase the χ^2^ statistic of a SNP more than LD to other categories:

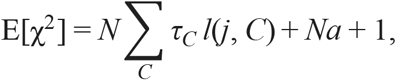

where *N* is the sample size, *τ*_*C*_ is the per-SNP heritability contributed by category *C, l*(*j, C*) is the LD score of SNP *j* to category *C*, defined as the summation of correlations with SNPs in category 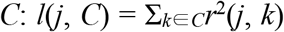 and *a* measures the contribution of confounding biases. We estimate *τ*_*C*_ for cis and trans separately by annotating *cis*- and *trans*-SNPs for each gene/protein using the 53 baseline functional categories^73^ (Finucane et al., 2015). The estimation of *cis* and *trans h*^2^ for each individual gene/protein has a large variance, therefore we applied constrained-intercept LD-score regression by fixing *a* at 0 and calculated *h*^2^ using the average estimated *τ*_*C*_ across all genes/proteins^18^:

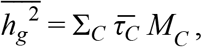

where 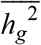 is the average total 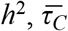 is the average estimated *τ*_*C*_ across all genes/proteins, and *M*_*C*_ is the total number of SNPs in category *C*.

#### Colocalization

Colocalization analyses are performed between pQTLs and 50 GWAS studies, including 29 blood traits, 8 autoimmune diseases, and 13 other traits, using the R package ‘coloc’ (v.5.1.0.1)^22^. For each pQTL as defined in the “pQTL data” section, colocalization analysis is performed with the corresponding locus in each of the GWAS studies. We only keep the GWAS loci with p value < 5×10^−8^, and set a posterior probability of hypothesis four (PP4) > 0.75 as the threshold for colocalization.

#### Bootstrapping and relevant p values

For the analyses comparing pLI score, proportion of heritability explained by trans, and enrichment of trans-pQTLs in PPI and TF, we first generate 1,000 bootstrapped samples using the R package ‘boot’ (v.1.3.28.1)^74^, then use the differences between bootstrapped samples to calculate an empirical p value:

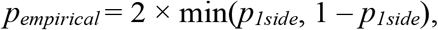

where *p*_*1side*_ is the proportion of the differences > 0 across 1,000 bootstrapped samples.

#### Estimation of QTL sharing using π_**1**_

To obtain a discovery set for eQTLs and pQTLs, we used genome-wide Bonferroni corrected p-values to obtain QTLs for each dataset. We estimated replication rates by using the Storey π_1_ statistics^24^ (Storey et al., 2003) on the set of p-values ascertained in the replication datasets (UKB-PPP for eQTLs and DGN/eQTLGen for pQTLs) of the most significant SNP-gene pairs in the discovery set. For all the leading SNPs of QTLs, we find the corresponding p values of SNP-gene/protein pairs in the replication set to calculate π_1_ using the R package ‘qvalue’ (v.2.24.0)^24^.

#### Enrichment of *trans*-pQTLs in PPI and TF

For each *trans-*pQTL from UKB-PPP, we considered the nearest protein-coding gene. To calculate the enrichment of those genes in TF, we first counted the number of those genes that are within the list of 1,639 putative human transcription factors from ^75^. We then randomly sampled a protein-coding gene with matched gene length for each of the nearest gene (length difference < 10% of the nearest gene) and counted the number of the randomly sampled genes that are within the TF list. We repeated the sampling 1,000 times to generate a random background and calculated the enrichment in TF as the ratio of the count between the nearest genes and random background.

To calculate the enrichment of nearest genes in PPI, we paired the nearest gene and the target gene of each *trans*-pQTL and counted the number of those pairs within PPIs from any of four databases: BioPlex^28^, CORUM^29^, HIPPIE^30^ and STRING^31^. We then randomly paired the nearest gene with a gene from all *trans*-pQTL target genes and counted the number of random pairs that are in PPI. We repeated the random pairing 1,000 times to generate a random background and calculated the enrichment in PPI as the ratio of the count between the observed pairs and random background. For *trans*-pQTLs of PPI clusters identified by *trans*-PCO, the calculation of enrichment of trans-pQTLs in PPI and TF is similar; however, for the enrichment in PPI, instead of investigating if the nearest gene and the target gene are in PPI, we investigated if the nearest gene is a subunit of the target PPI cluster.

#### QTL annotation

For the annotation of *trans*-eQTLs and *trans*-pQTLs, we used the leading SNP of each *trans*-QTL and SnpEff (v.5.2c)^34^ to annotate those SNPs. The proportion of each annotation for *trans*-eQTL and *trans*-pQTL was compared using Fisher’s exact test.

#### Enrichment of *trans*-pQTLs at PPI interface

Genomic regions of PPI interfaces are obtained from INCIDER^35^. Enrichment of *trans*-pQTL and *trans*-eQTL at PPI interfaces is calculated as follows. For the leading SNP of each *trans*-QTL, we randomly selected a coding SNP from the UKB-PPP genotype for 1,000 times. We matched the distance to TSS of the nearest protein coding gene and minor allele frequency (MAF) between leading SNPs and randomly selected SNPs, i.e., the differences of the distance and MAF between the leading SNP and the selected SNP have to be smaller than 10% to avoid the bias caused by distance to TSS and MAF. Then the enrichment is calculated as the ratio between the proportion of leading SNPs of *trans*-QTLs and the proportion of randomly selected SNPs located at PPI interfaces.

#### Protein clustering based on PPI

We defined protein clusters based on PPI from BioPlex^28^, CORUM^29^, HIPPIE^30^ and STRING^31^. CORUM is a well-curated protein complex database, thus we used each protein complex directly as a protein cluster. STRING calculates the pairwise protein–protein association score based on PPIs from multiple sources. We recalculated the score based on only PPIs with strong evidence, i.e., those from curated databases or experimentally determined, and kept the protein pairs with association scores > 0.75. We generated PPI clusters using the MCL algorithm^36^ for pairwise PPIs in BioPlex, HIPPIE, and STRING separately. We ran the MCL algorithm with a default inflation value equal to 2. We then merged the protein clusters from 4 databases and retained the proteins that overlapped with those measured in UKB-PPP. After removing repeated clusters and sub-clusters, we obtained 1,088 clusters comprising 2,165 unique proteins.

#### Identify trans-pQTLs using trans-PCO

We use trans-PCO to identify trans-pQTLs of PPI clusters based on summary statistics^8^. For a PPI cluster with *K* proteins:

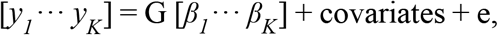

where *y*_*k*_ is the expression level of the kth protein for *k* = 1, …, *K*; *G* is the SNP genotype, *β*_*k*_ is the genetic effect of the SNP on the *k*-th protein, covariates are the fixed effects, and *e* is the residual. The null hypothesis is that all the genetic effects are 0:

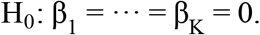

A single PC-based test to test this null hypothesis is:

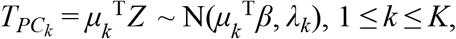

where *Z* is a *K*×1 vector of univariate z-scores of the SNP for *K* proteins, *μ*_*k*_ is the *k*-th eigenvector of the correlation matrix Σ of *Z, β* is the vector genetic effects of the SNP on *K* proteins, *λ*_*k*_ is the *k*-th eigenvalue of Σ. The test statistic follows a normal distribution under the null hypothesis, and we denote the p value of the test as *p*_*k*_.

A single PC-based test has only limited power, while PC-based omnibus test (PCO) combines multiple single PC-based tests in linear and nonlinear ways to achieve higher power^76^. There are six tests included in PCO^76^:

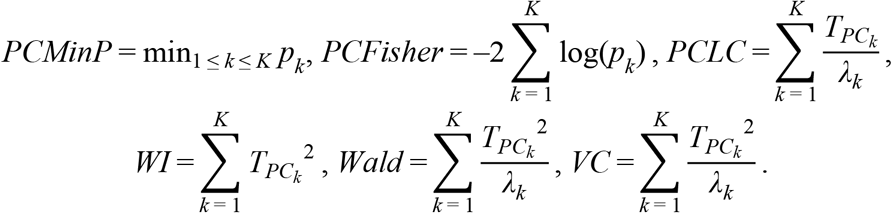

Then we use ACAT^77^ to combine the p values of the six tests:

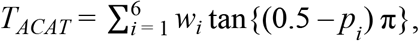

where *w*_*i*_ is the weight of each test which is set to be equal (1/6), and *p*_*i*_ is the p value of the *i*-th test. The test statistic follows a Cauchy distribution under the null hypothesis.

The inputs of trans-PCO include z-scores of SNP-protein pairs and the correlation matrix Σ of *Z* for each PPI cluster. The correlation matrix Σ can be estimated either using the protein expression profile or insignificant z-scores^8^. Since only summary statistics are available from UKB-PPP, we estimate Σ using insignificant z-scores. Specifically, we select independent SNPs that are insignificantly associated (univariate p > 1×10^−4^) with all proteins in the PPI cluster, and then calculate the sample correlation matrix using the z-scores of proteins over these independent null SNPs. For a SNP and PPI cluster pair, *trans*-PCO only keeps the proteins that are encoded on a different chromosome than the test SNP to limit the analysis to *trans* associations.

#### Functional enrichment of PPIs

We use the web server g:Profiler^78^ to run functional enrichment analysis for genes of each PPI. We include three Gene Ontology (GO) term groups, biological process (BP), molecular function (MF), and cellular component (CC) in the analysis. g:Profiler uses Fisher’s one-tailed test to measure the enrichment of ontology terms. A Benjamini-Hochberg FDR < 0.05 is used as the threshold to decide a significant GO term, and the results are reported as the FDR adjusted p values.

## Acknowledgements

We thank N. Gonzales and S. Sumner for editing the manuscript. This work was completed in part with resources provided by the University of Chicago’s Research Computing Center. This research was funded by two NIGMS Maximizing Investigators’ Research Awards (R35GM138084 to X.L and R35GM153249 to Y.I.L.).

## Author contributions

L.J., Y.I.L., and X.L. designed analyses. L.J. performed all data analysis under the supervision of X.L.. L.J., Y.I.L., and X.L. wrote the manuscript.

