## Supplementary figures for "Protein–protein interactions shape *trans*-regulatory impact of genetic variation on protein expression and complex traits"

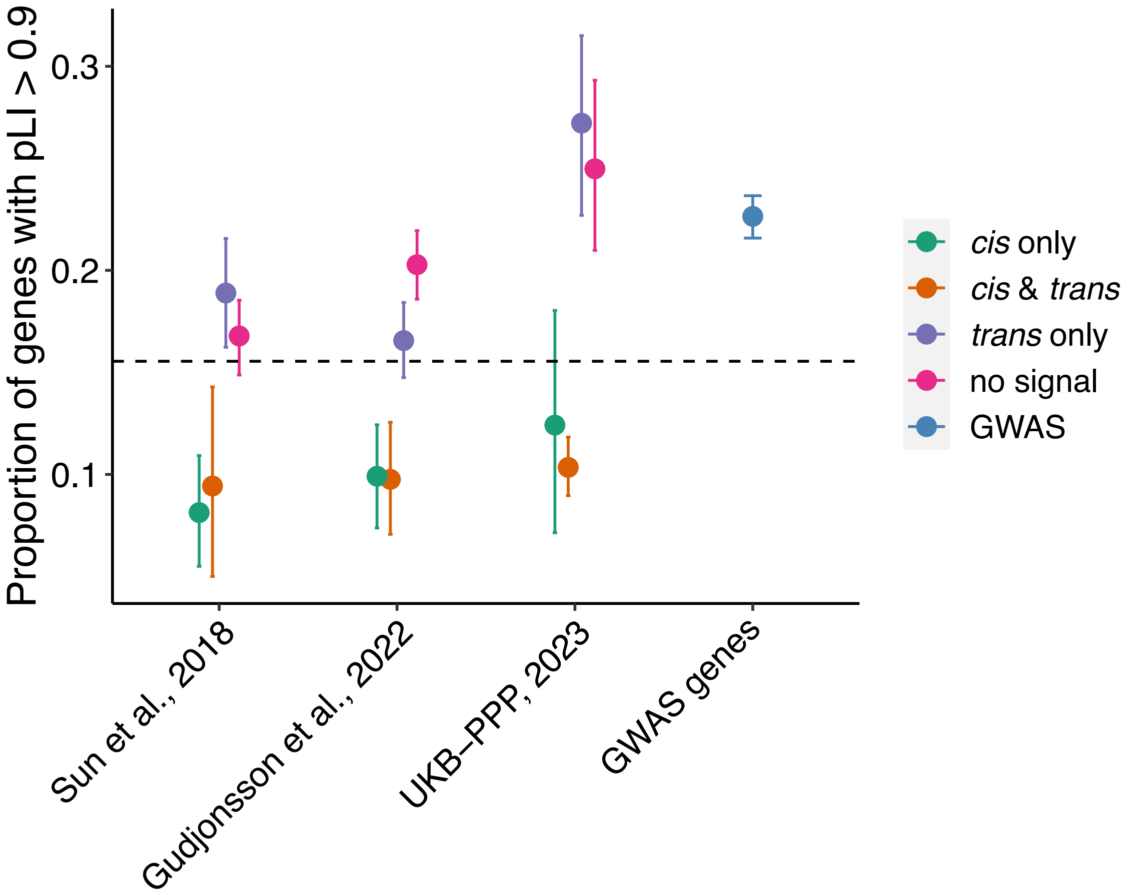


Figure S1 Proportion of pQTL target genes with pLI score > 0.9 three plasmatic proteome datasets[1–3] and GWAS genes reported by [4]. Dashed line shows the proportion of high-pLI genes across the whole genome. Genes are categorized into 4 groups: genes with both *cis*- and *trans*-pQTLs; with *cis*-pQTLs only; with *trans*-pQTLs only; and with no significant pQTLs.

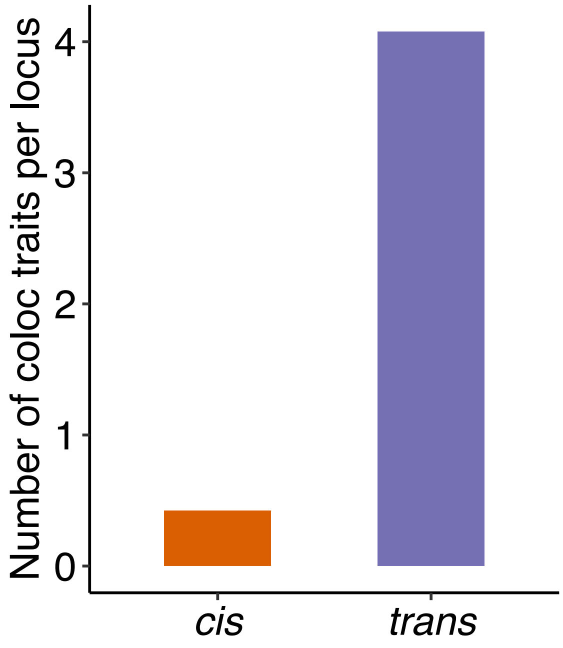


Figure S2 Average number of colocalized traits per cis- and *trans*-pQTL from UKB-PPP.

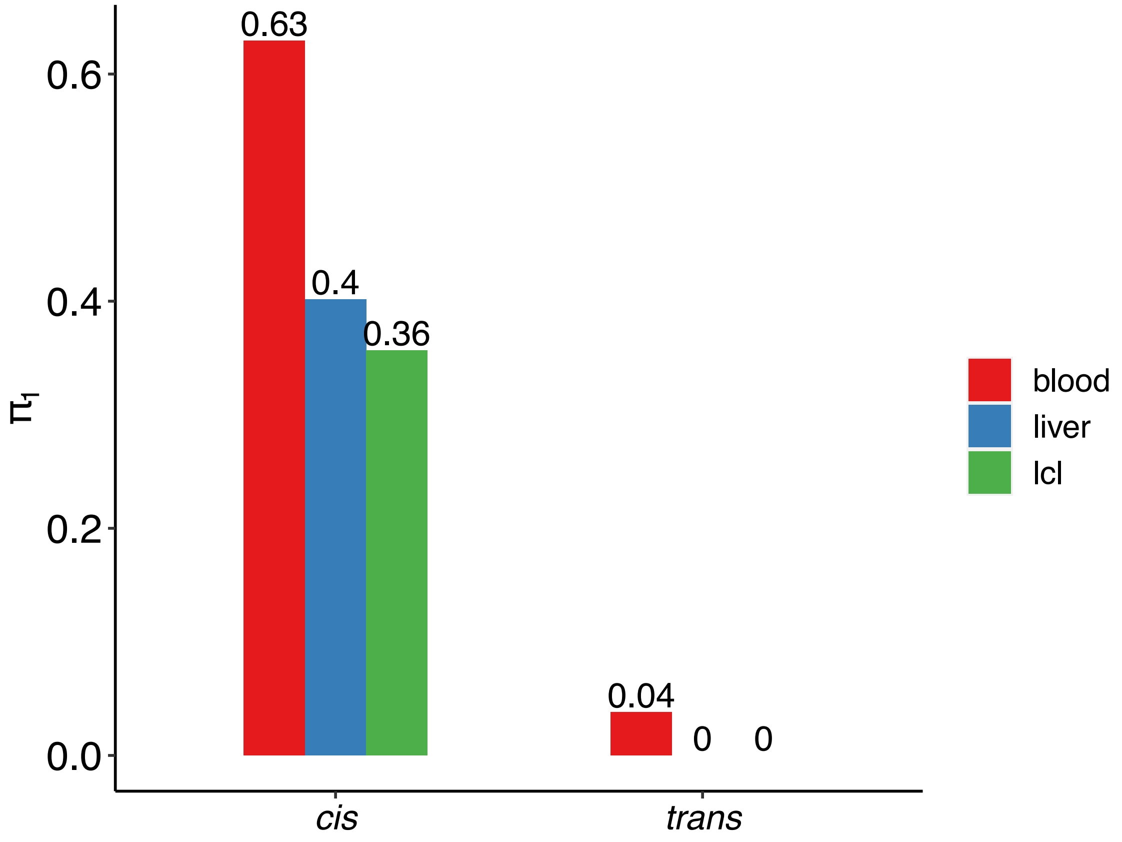


Figure S3 Estimated proportion of significant signals ($\pi_1$) for pQTLs of UKB-PPP in three GTEx tissues.

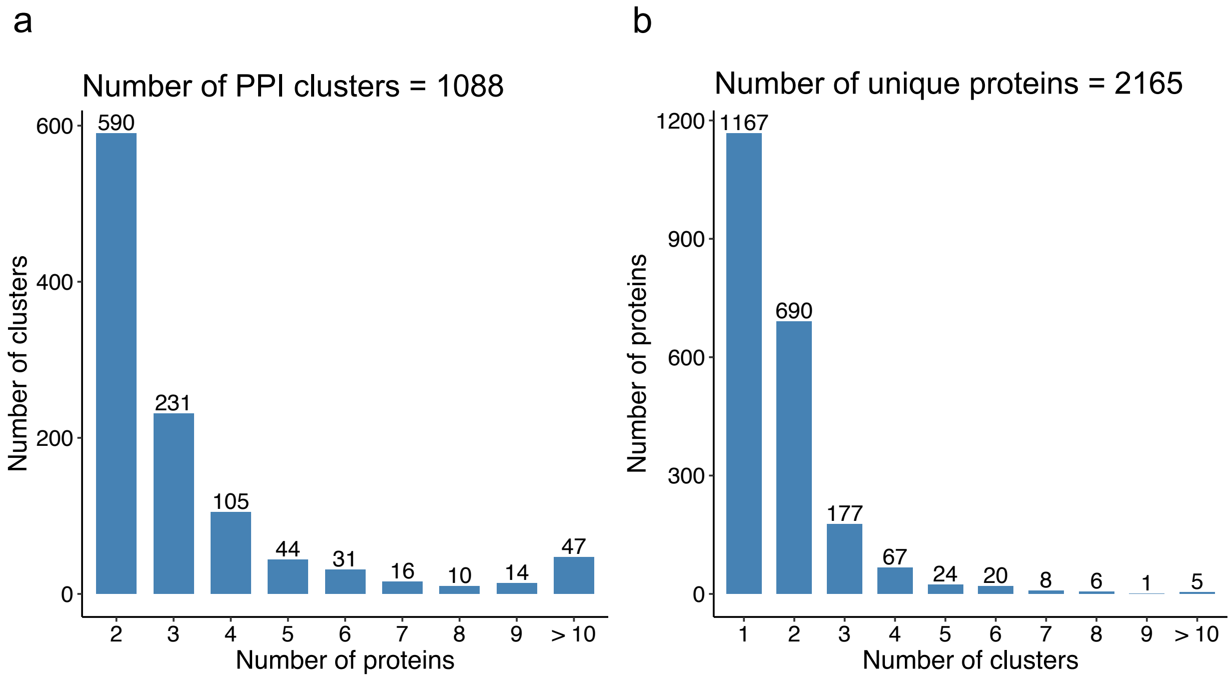


Figure S4 Summary of PPI clusters generated based on 4 PPI databases, including BioPlex[5], HIPPIE[6], STRING[7], and CORUM[8].

a. Histogram of the number of proteins per cluster.

b. Histogram of the number of clusters each protein belongs to.

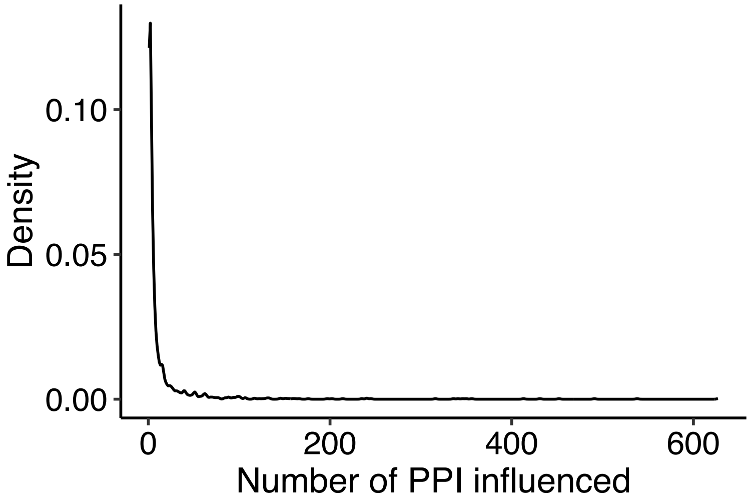


Figure S5 Distribution of number of PPI influenced by each independent *trans*-pQTL locus.

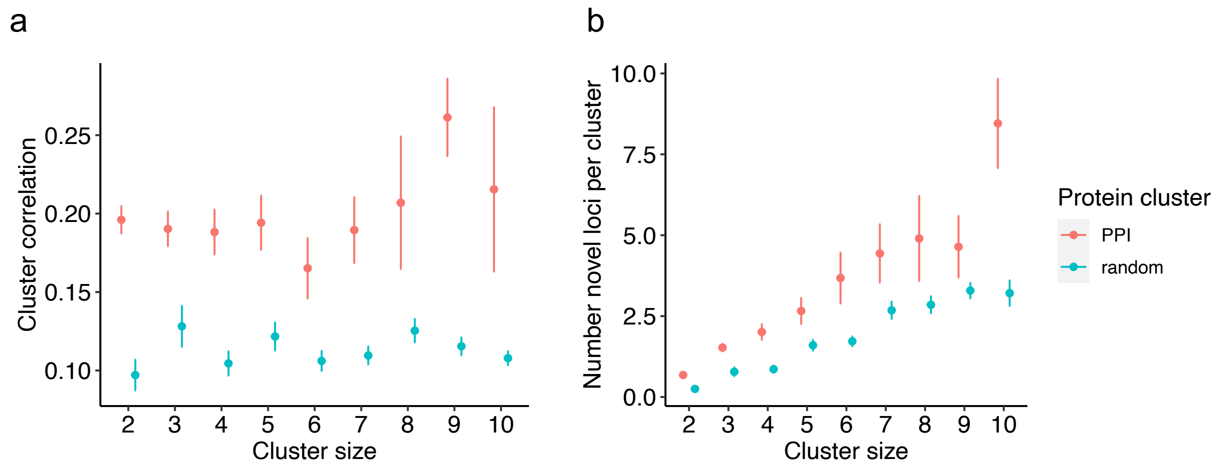


Figure S6 Comparison between PPI protein clusters and random clusters across cluster size from 2 to 10. Each cluster size includes 100 randomly generated protein clusters. The error bar represents the standard error.

a. Mean of the absolute pairwise correlations for PPI clusters and random clusters.

b. Number of novel *trans*-pQTLs for PPI clusters and random clusters.

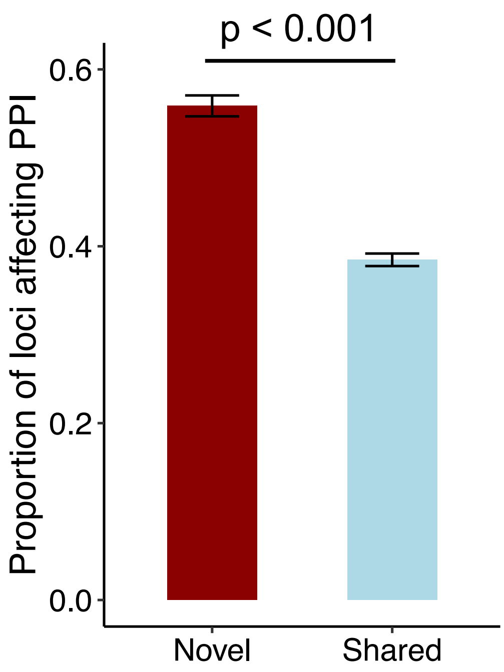


Figure S7 Proportion of novel and shared *trans*-pQTLs having multiple univariate associations in the PPI.

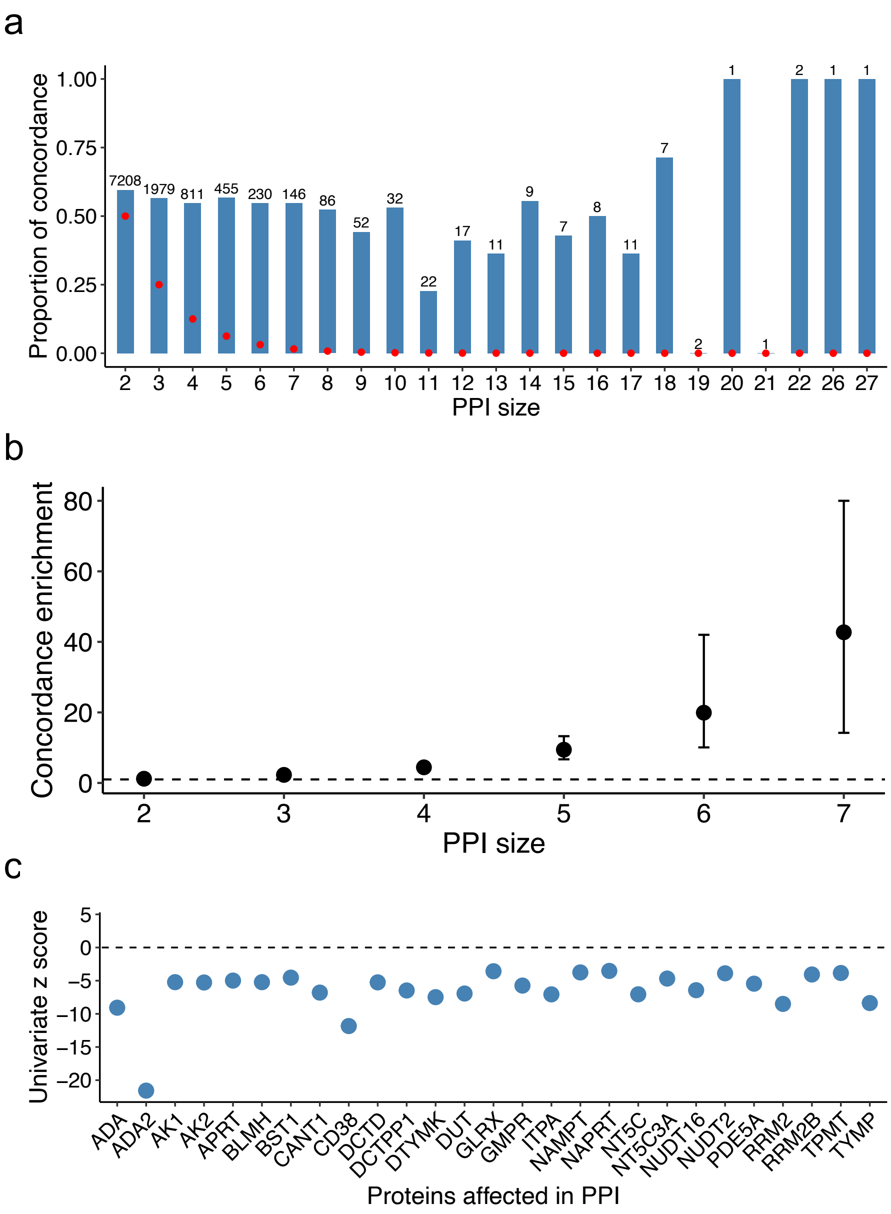


Figure S8 Effect direction concordance of *trans*-pQTLs affecting multiple proteins (univariate p < 0.05/|PPI|) in the target PPI.

a. Proportion of *trans*-pQTLs having the same effect direction on all proteins of the same PPI across different PPI sizes. Numbers of *trans*-pQTLs investigated for each PPI size are marked above each bar. The red dots represent the expected proportion of concordance when each effect has random directions.

b. Enrichment of *trans*-pQTLs having concordant effect for PPI sizes from 2 to 7. Enrichment is calculated as the ratio between proportion of *trans*-pQTLs having concordant effect for observed effect directions and randomly generated effect directions. Error bars are 95% confidence interval based on 1,000 random samples. Random samples have very few *trans*-pQTLs with concordant effect for PPI size > 8, thus are not shown in the figure.

c. Example of a *trans*-pQTL of PPI cluster 953 having negative effect on all 27 affected proteins.
